# The microRNA, *miR-18a*, regulates NeuroD and photoreceptor differentiation in the retina of the zebrafish

**DOI:** 10.1101/440263

**Authors:** Scott M. Taylor, Emily Giuffre, Patience Moseley, Peter F. Hitchcock

## Abstract

During embryonic retinal development, six types of retinal neurons are generated from a pool of multipotent progenitors in a strict spatiotemporal pattern. This pattern requires cell cycle exit (i.e. neurogenesis) and differentiation to be precisely regulated in a lineage-specific manner. In zebrafish, the bHLH transcription factor NeuroD governs photoreceptor genesis through Notch signaling but also governs photoreceptor differentiation though distinct mechanisms that are currently unknown. Also unknown are the mechanisms that regulate NeuroD and the spatiotemporal pattern of photoreceptor development. Members of the *miR-17-92* microRNA cluster regulate CNS neurogenesis, and a member of this cluster, *miR-18a*, is predicted to target *neuroD* mRNA. The purpose of this study was to determine if *miR-18a* regulates NeuroD in the retina and if it plays a role in photoreceptor development. Quantitative RT-PCR showed that, of the three *miR-18* family members (*miR-18a*, *b* and *c*), *miR-18a* expression most closely parallels *neuroD* expression. Morpholino oligonucleotides and CRISPR/Cas9 gene editing were used for *miR-18a* loss-of-function (LOF) and both approaches resulted in larvae with more mature photoreceptors at 70 hpf without affecting cell proliferation. Western blot showed that *miR-18a* LOF increases NeuroD protein levels and *in vitro* dual luciferase assay showed that *miR-18a* directly interacts with the 32UTR of *neuroD*. Finally, *tgif1* mutants have increased *miR-18a* expression, less NeuroD protein and fewer mature photoreceptors, and the photoreceptor deficiency is rescued by *miR-18a* knockdown. Together these results show that, independent of neurogenesis, *miR-18a* regulates the timing of photoreceptor differentiation and indicate that this occurs through post-transcriptional regulation of NeuroD.

## INTRODUCTION

In the developing retina, six types of neurons are generated from a pool of multipotent, mitotic progenitors in a sequence that is highly conserved among vertebrates (Bassett & Wallace, 2012; Centanin & Wittbrodt, 2014; Wallace, 2011). For mature neurons to develop, progenitors must be specified to a particular fate, exit the cell cycle, and differentiate into mature neurons. These events are governed (in part) by transcription factors that regulate expression of genes involved in the cell cycle and neuronal differentiation. The basic-loop-helix (bHLH) transcription factors play prominent roles in these events (Akagi et al., 2004; Baker & Brown, 2018; Joseph A. Brzezinski, Kim, Johnson, & Reh, 2011; Mao et al., 2013; Ohsawa & Kageyama, 2008; Pollak et al., 2013). Rod and cone photoreceptors are the neurons in the distal retinal layer that first collect visual information and, in zebrafish, the bHLH transcription factor NeuroD governs the cell cycle in photoreceptor progenitors through intercellular Notch signaling (Malgorzata J. Ochocinska & Hitchcock, 2007; Taylor et al., 2015). Following cell cycle exit, NeuroD also governs photoreceptor differentiation through separate mechanisms that are currently unknown.

In the embryonic zebrafish retina, *neuroD* mRNA is expressed from 30 hours postfertilization (hpf) and, by 48 hpf, is expressed in all photoreceptor progenitors in the developing outer nuclear layer (ONL) (Malgorzata J. Ochocinska & Hitchcock, 2007). Most photoreceptor genesis and differentiation occur between 48 and 72 hpf beginning in a small ventronasal region called the precocious ventral patch (Schmitt & Dowling, 1999), then spreading peripherally throughout the ONL with cones differentiating slightly before rods (Stenkamp, 2007). This tightly controlled spatiotemporal pattern of photoreceptor differentiation, despite the constitutive expression of *neuroD* throughout the ONL, suggests that post-transcriptional mechanisms may regulate NeuroD and the timing of photoreceptor differentiation.

Post-transcriptional regulation can occur through small ~22 nucleotide (nt) single stranded RNA molecules called microRNAs (miRNAS) that bind to target mRNA through complementary base paring and regulate protein expression by blocking translation and/or causing mRNA degradation (Huntzinger & Izaurralde, 2011). Several miRNAs have been shown to regulate key aspects of brain and retinal development (Andreeva & Cooper, 2014; La Torre, Georgi, & Reh, 2013; Madelaine et al., 2017; Ohana et al., 2015; Petri, Malmevik, Fasching, Åkerblom, & Jakobsson, 2014; Sundermeier & Palczewski, 2016) and miRNAs are investigated here as potential regulators of NeuroD and photoreceptor genesis. MicroRNAs are initially expressed as primary transcripts called pri-miRNAs, are then cleaved by the Drosha enzyme into shorter precursors (pre-miRNAs) that fold into imperfect stem-loop structures and are ultimately cleaved in the cytoplasm by Dicer to become mature miRNAs (Winter, Jung, Keller, Gregory, & Diederichs, 2009; Zeng, Yi, & Cullen, 2005). Mature miRNAs typically function by binding via a specific “seed” sequence comprising ~6-8 nucleotides near the 5’ end of the miRNA, to a complimentary sequence in the 3’ untranslated region (UTR) of the target mRNA (Bartel, 2004; Broughton, Lovci, Huang, Yeo, & Pasquinelli, 2016). Based on these complimentary sequences, interactions between miRNAs and target mRNAs can be predicted (e.g. www.targetscan.org). A single miRNA can potentially regulate hundreds of different mRNAs and a single mRNA can be targeted by many different miRNAs (Peter, 2010), making it difficult to identify functional relationships between miRNAs and specific targets. Understanding the functions of miRNAs might, therefore, require combined approaches using morpholinos or siRNAs that block multiple functionally overlapping miRNAs (Alex Sutton Flynt, Rao, & Patton, 2017), as well as gene editing technologies (e.g. CRISPR/Cas9, TALENS) that disrupt individual miRNAs by generating insertion/deletion (indel) mutations in miRNA genes.

Many miRNAs are transcribed together as polycistronic clusters that are processed into functionally distinct miRNAs (Khuu, Utheim, & Sehic, 2016). The *miR-17-92* cluster generates 15 mature miRNAs including *miR-19b*, which regulates NeuroD and insulin secretion in the pancreas (Zhang et al., 2011), and several miRNAs that regulate neurogenesis in the mouse neocortex (Bian et al., 2013). Another member of this cluster, *miR-18a*, is also predicted to interact with *neuroD* (www.targetscan.org/fish_62/) but has not been studied in the developing brain or retina. Additionally, two other members of the *miR-18* subfamily, *miR-18b* and *miR-18c*, are at distinct genetic loci and not part of the *miR-17-92* cluster but have identical seed sequences to *miR-18a* and are also predicted to target *neuroD*.

Based on their predicted interactions with *neuroD*, *miR-18a*, *b and c* were examined as potential post-transcriptional regulators of NeuroD during embryonic photoreceptor genesis. Quantitative PCR (qPCR) showed that, of the three miRNAs, the timing of *pre-miR-18a* and *miR-18a* expression most closely parallel that of *neuroD.* Morpholino oligonucleotides targeted to *miR-18a*, *b* or *c* produced an identical phenotype with increased numbers of photoreceptors at 70 hpf. Focusing solely on *miR-18a*, an in-vitro dual luciferase assay showed that *miR-18a* interacts directly with the 3’ UTR of *neuroD* mRNA. Mutation of *miR-18a* using CRISPR/Cas9 gene editing reproduced the morphant phenotype, where more mature rod and cone photoreceptors are present at 70 hpf with no effect on cell proliferation. Western blot showed that when photoreceptor differentiation begins at 48 hpf, knockdown or mutation of *miR-18a* results in higher levels of NeuroD protein. Finally, in *tgif1* mutant embryos that have higher levels of *miR-18a*, there is less NeuroD protein and fewer mature photoreceptors, and the photoreceptor deficiency is rescued by *miR-18a* knockdown. Taken together, these data show that during embryonic development, *miR-18a* regulates the timing of differentiation in post-mitotic photoreceptors and indicate that *miR-18a* functions through post-transcriptional regulation of NeuroD.

## METHODS

### PCR Methods

AB wild type (WT) strain zebrafish, purchased from the Zebrafish International Research Center (ZIRC; University of Oregon, Portland, OR, USA), were used for developmental experiments and to generate *miR-18a* mutants. Embryos were collected within 15 minutes of spawning and incubated at 28.5°C on a 14/10-hour light/dark cycle. For standard qPCR used to amplify *miR-18a, b* and *c* precursor molecules, total RNA was collected from 40 whole embryo heads or 40 whole eyes (at 70 hpf) per biological replicate, using the Aurum Total RNA Mini Kit and following the manufacturer’s protocol (Bio-Rad Laboratories, Inc., Hercules, CA, USA). Reverse transcription was performed using the Qiagen QuantiTect Reverse Transcription Kit by following the manufacturer’s protocol (Qiagen, Venlo, The Netherlands). Forward and reverse primers used to amplify *miR-18a, b* or *c* precursor sequences were as follows: *pre-miR-18a* F:GGCTTTGTGCTAAGGTGCATCTAG; R:CAGAAGGAGCACTTAGGGCAGTAG; *pre-miR-18b* F:CTGCTTATGCTAAGGTGCATTTAG; R:CTTATGCCAGAAGGGGCACTTAGG; *pre-miR-18c* F:GCCTTCCTGCTAAGGTGCATCTTG; R:CCTGCCAAAAGGAACATCTAGCGC. The primers used for qPCR analysis of *neuroD* mRNA expression were F:ATGCTGGAGTCTCAGAGCAGCTCG; R:AACTTTGCGCAGGCTCTCAAGCGC. Biological qPCR replicates were each performed in triplicate using 20 ng cDNA and IQ SYBR Green Supermix (Bio-Rad Laboratories, Inc.) and run on a Bio-Rad 384-well real-time PCR machine. Relative fold changes in expression levels were calculated using the comparative C_T_ method and, when applicable, were compared for statistical significance using a Student’s t-test with a significance level of p<0.05.

For qPCR analysis of mature *miR-18a* expression, a TaqMan custom qPCR assay was designed for mature *miR-18a* and for the small nuclear RNA *U6*, to be used as the housekeeping gene for data normalization (ThermoFisher Scientific, Halethorp, MD, USA). Total RNA, including small RNAs, were collected using a mirVana miRNA isolation kit (AM1560; ThermoFisher Scientific). For comparison of precursor and mature miR expression, using the same samples, standard reverse transcription and qPCR were performed for *pre-miR-18a* amplification as described above and primer-specific TaqMan reverse transcription and mature miRNA qPCR were performed using the manufacturer’s protocol (ThermoFisher Scientific). For mature miRNA qPCR, *miR-18a* expression was normalized to *U6* expression, relative to the 30 hpf sample, and was calculated using the comparative C_T_ method.

### miRNA Knockdown with Morpholino Oligonucleotides

Morpholino oligonucleotides (MO; Gene Tools, LLC, Philomath, OR. USA) targeted to the mature strand of *miR-18a*, *miR-18b* or *miR-18c* were used to induce *miRNA* knockdown. The *miR-18a* morpholino [5‘-CTATCTGCACTAGATGCACCTTAG-3’] was published previously and shown to effectively knock down *miR-18a* in vivo (Friedman et al., 2009). The *miR-18b* [5‘-CTATCTGCACTA<u>A</u>ATGCACCTTAG-3’] MO used here differs from the *miR-18a* MO by only one nucleotide (underlined) and the *miR-18c* MO [5‘-CTA<u>A</u>CT<u>A</u>CAC<u>A</u>AGATGCACCTTAG-3’] differs by only three nucleotides. Morpholino oligonucleotides were diluted in 1X Daneau buffer (Nasevicius & Ekker, 2000), and 3 ng MO were injected at the single cell stage as described previously (M. J. Ochocinska & Hitchcock, 2009).

### Systemic Labeling with 5-Bromo-2’-Deoxyuridine (BrdU), Immunohistochemistry and Cell Counting

Cells in S-phase of the cell cycle were labeled by incubating embryos for 20 minutes, immediately prior to sacrifice, in ice-cold 10 mM BrdU dissolved in embryo rearing solution containing 15% dimethylsulfoxide (DMSO). Whole embryos were fixed and prepared for histology as previously described (Taylor et al., 2015), embedded in optical cutting temperature (OCT) medium, and heads were sectioned at 10 μm and mounted on glass slides (Superfrost plus; Fisher Scientific, Pittsburgh, PA). Immunolabeling was performed using previously published protocols (Luo et al., 2012) on cross-sections through the central retina in the vicinity of the optic nerve. For BrdU immunolabeling, DNA was denatured by incubating sections in 100°C sodium citrate buffer (10mM sodium citrate, 0.05% Tween 20, pH 6.0) for 30 minutes, cooled at room temperature for 20 minutes, and processed with standard immunolabeling techniques. The primary and secondary antibodies and dilution factors used here were: mouse anti-BrdU, 1:100 (347580; BD Biosciences, Franklin Lakes, NJ, USA); Zpr-1,1:200 (anti-Arrestin 3, red-green double cones, ZIRC); goat anti-mouse Alexa Fluor 488 and goat anti-mouse Alexa Fluor 555, 1:500 (Life Technologies, Carlsbad, CA, USA). Nuclei were counterstained with 20 mM Hoechst 33342 (ThermoFisher Scientific) prior to adding coverslips. BrdU-labeled cells and cones were counted in the central-most cross-section for each larval fish. Cell counts were compared using a Student’s t-test, with a p<0.05 indicating statistically-significant differences.

### Luciferase assay

To test the interaction between *miR-18a* and the 3’ UTR of *neuroD* mRNA, an in vitro dual luciferase assay was performed following published protocols (Jin, Chen, Liu, & Zhou,2012). Briefly, a custom oligonucleotide corresponding to a 66 bp portion of the *neuroD* 3’ UTR containing the predicted target sequence for *miR-18a* (underlined) *[“neuroDWT”* 5‘-GGAGAAAAGAGAATTGGTTGATTCTCGTT<u>CACCTTA</u>TGT ATTGTATTCTATAGCGCTTCTACGTT G-3’] was generated (ThermoFisher Scientific) and inserted into the pGL3 vector immediately 3’ of the firefly luciferase gene (E1741; Promega, Madison, WI, USA). A negative control pGL3 vector was also created containing the same *neuroD* 3’ UTR sequence, but with the predicted *miR-18a* target site mutated to <u>TTTTTTT</u> [*“neuroDMut”*]. Following bacterial transformation, culture and plasmid purification, HEK 293 cells, grown to 20-40% confluence, were transfected with either the *neuroDWT* or *neuroDMut* plasmid along with the pRL-TK vector that constitutively expresses *Renilla* luciferase to serve as an internal transformation control. Each transfection group was co-transfected with either *hsa-miR-18a-5p* mimic (identical to zebrafish mature *miR-18a*) or *hsa-let7a-5p* mimic, for which neither vector had a predicted target site and served as a negative control (Exiqon/Qiagen, Venlo, The Netherlands). Following transfection, 48 h incubation and cell lysis, luciferase expression levels were assayed on a luminometer using a Dual-Luciferase Reporter Assay System (E1910; Promega). Using this approach, direct interaction between the miRNA mimic and the cloned 3’ UTR *neuroD* sequence is expected to reduce the level of firefly luciferase levels. For each experimental group, firefly luciferase was normalized to constitutive *Renilla* luciferase levels and results were compared using a Student’s t-test.

### Generating *miR-18a* mutants

CRISPR/Cas9 genome editing was used to generate mutations in the *miR-18a* gene (Taylor et al., 2015). Briefly, the sgRNA target sequence was identified within the *miR-18a* precursor sequence using ZiFiT software (available in the public domain at www.zifit.partners/org). Cas9 mRNA and sgRNA were generated (Hwang et al., 2013), and single cell-stage embryos were injected with 1 nL solution containing 100 pg/nL sg RNA and 150 pg/nl Cas9 mRNA. F0 injected fish were raised to maturity. Genomic DNA was purified from caudal fins, and screening primers (F: CCAGGAAAGATGGGAGTAGTTG; R: CTCACACTGCAGTAGATGACAG) were used to amplify a 626 bp region around the sgRNA target site using standard PCR and 100 ng template DNA. CRISPR-induced insertions and deletions were detected using the T7 endonuclease assay according to established protocols (available in the public domain at www.crisprflydesign.org). Briefly, 200ng of purified PCR product was used for the analysis and, following denaturation and reannealing, subjected to a 15-minute digest at 37°C with 10 U T7 endonuclease I (New England Biolabs, Ipswich, MA, USA). Digested DNA was run on a 2% agarose gel and indels were identified by the presence of a double band around 200-300 bp. F0 adult fish that were positive for indels were outcrossed with AB WT fish and, using the same methods as above, the T7 assay was used to identify indels in F1 generation adults. PCR products from T7-positive F1 adults were subcloned into the pGEM-T Easy vector (Promega) and 6 clones were sequenced for each fish. Mutations were detected using a pairwise blast (NCBI, Bethesda, DM, USA) against WT DNA.

In one F1 adult fish, a 25 bp insertion was detected in the *miR-18a* precursor sequence, and this introduced an AleI restriction enzyme cut site that was used for subsequent genotyping. The same screening primers (above) were used for this method and AleI digest using standard protocols (New England Biolabs) cuts the mutant PCR product into 244 bp and 407 bp segments that are easily visualized using gel electrophoresis. F1 generation heterozygous adult fish were out-crossed and then F2 generation heterozygotes in-crossed to produce a homozygous line of *miR-18a* mutants, and these fish were in-crossed to produce homozygous mutant embryos used here.

### Western blot analysis

Protein samples were obtained by pooling whole heads of embryos or larvae at 48 or 70 hpf, respectively, in RIPA lysis buffer (89900; ThermoFisher Scientific) containing 1x protease and phosphatase inhibitor cocktail (5872; Cell Signaling Technology, Danvers, MA, USA). Proteins were separated in a 12% SDS-PAGE pre-cast gel (4561043; Bio-Rad Laboratories, Inc.) and transferred to a PVDF membrane (Sigma-Aldrich Corp., St. Louis, MO, USA). The membrane was incubated in 5% bovine serum albumin (BSA) with 0.05% Tween-20 for 2 hours to block non-specific binding of the antibodies and then incubated overnight at 4C with rabbit anti-NeuroD antibodies (M.J. Ochocinska & Hitchcock, 2009) diluted 1:1000 in 2.5% blocking solution. Blots were rinsed with TBS with 0.05% Tween-20 and incubated with horseradish peroxidase-conjugated secondary IgG (1:2000) for 1 hour at room temperature. Bands were visualized using the enhanced chemiluminescence assay detection system (34075; ThermoFisher Scientific). For loading controls, blots were stripped in stripping buffer (21059; ThermoFisher Scientific) for 5 minutes, processed as described above and labeled with mouse anti-βactin antibodies (1:5000) (NB10074340T, Novus Biologicals, LLC, Littleton, CO, USA). Images were captured using the FluorChem E Imaging System (Bio-Techne, Minneapolis, MN, USA) and band intensity was quantified relative to βactin.

### In situ Hybridization

*In situ* hybridization with *rhodopsin* probes was used to identify rod photoreceptors. A DIG-labeled antisense riboprobe for zebrafish *rhodopsin* was generated from a 976 bp PCR product containing a T3 polymerase promoter sequence (lowercase, underlined) on the reverse primer <u>f aattaaccctcactaaaggg</u>CTTCGAAGGGGTTCTTGCCGC1 following published methods (David & Wedlich, 2001); for similar primer lengths, a T7 polymerase promoter sequence (lowercase, underlined) was added to the forward primer (<u>taatacgactcactataggg</u>GAGGGACCGGCATTCTACGTG). The antisense DIG-labeled probe was generated using T3 polymerase and in situ hybridization performed as previously described (Barthel & Raymond, 1993; Malgorzata J. Ochocinska & Hitchcock, 2007; Taylor et al., 2015). Control and morphant or mutant sections were mounted on the same slides and color reactions were developed for identical periods of time. Cells were counted in cross-sections and compared as described above.

For in situ hybridization labeling of *miR-18a* in tissue sections, a miRCURY LNA detection probe (Exiqon/Qiagen), labled with DIG at the 5’ and 3’ ends, was designed to hybridize with the mature *miR-18a* sequence. Standard in situ hybridization methods were used, as described above, using a 0.25μM probe working concentration at a hybridization temperature of 58°C.

## RESULTS

### *miR-18a* expression closely parallels *neuroD* expression

In the developing brain and retina, *neuroD* mRNA expression increases markedly between 30 and 72 hpf (Figure 1A), and most photoreceptors are generated between 48 and 72 hpf (Stenkamp, 2007). As a first step to determine if *miR-18* might regulate photoreceptor development, qPCR was used to analyze expression at time points consistent with *neuroD* and photoreceptor genesis. The miRNAs *miR-18a*, *mir-18b* and *mir-18c* differ in sequence by only 1-3 nucleotides, are expressed from distinct genetic loci, and have a conserved seed sequence that is predicted to interact with *neuroD* mRNA. Quantitative PCR analysis of mature miRNAs requires specialized kits (e.g. TaqMan, Applied Biosystems) and the nearly identical sequences among closely related miRNAs (e.g. miR-18a, b and c) can result in cross-amplification. The longer precursor molecules (pre-miRNAs) for similar miRNAs, however, have unique sequences and can be analyzed with standard qPCR and, if expression levels are proportional to mature miRNAs, can be used as a fast and accurate proxy for mature miRNA expression. To determine if *pre-miR-18a* expression can be used as a proxy for mature *miR-18a*, total RNA (including short RNAs) was purified from whole head tissue of zebrafish embryos at 30, 48 and 70 hpf with a miRVana miRNA isolation kit and then, on the same samples, standard qPCR was performed for *pre-miR-18a* and TaqMan qPCR for mature *miR-18a*. The results showed that the levels of both mature *pre-miR-18a* and *miR-18a* increase steadily and proportionally between 30 and 70 hpf (Figure 1B), indicating that *pre-miR-18a* can be used as a proxy for mature *miR-18a* expression. Standard qPCR was then used to compare expression of the 83-87 nt pre-miRNAs for *miR-18a*, *b* and *c* that each have unique sequences, despite the nearly identical mature miRNA sequences. In brain and retina tissue, identically to *neuroD* mRNA, *pre-miR-18a* expression increases steadily between 30 and 70 hpf, while *pre-miR-18b* expression remains substantially lower at all time points and *miR-18c* expression decreases after 30 hpf (Figure 1C). By 70 hpf, eyes are large enough for easy dissection and qPCR analysis of eye tissue only, and this showed that *pre-miR-18b* expression is substantially lower in the eye compared with *pre-miR-18a* or *c* (Figure 1D). These results show that among the three *pre-miRs, pre-miR-18a* expression most closely parallels that of *neuroD*.

**Figure 1.**
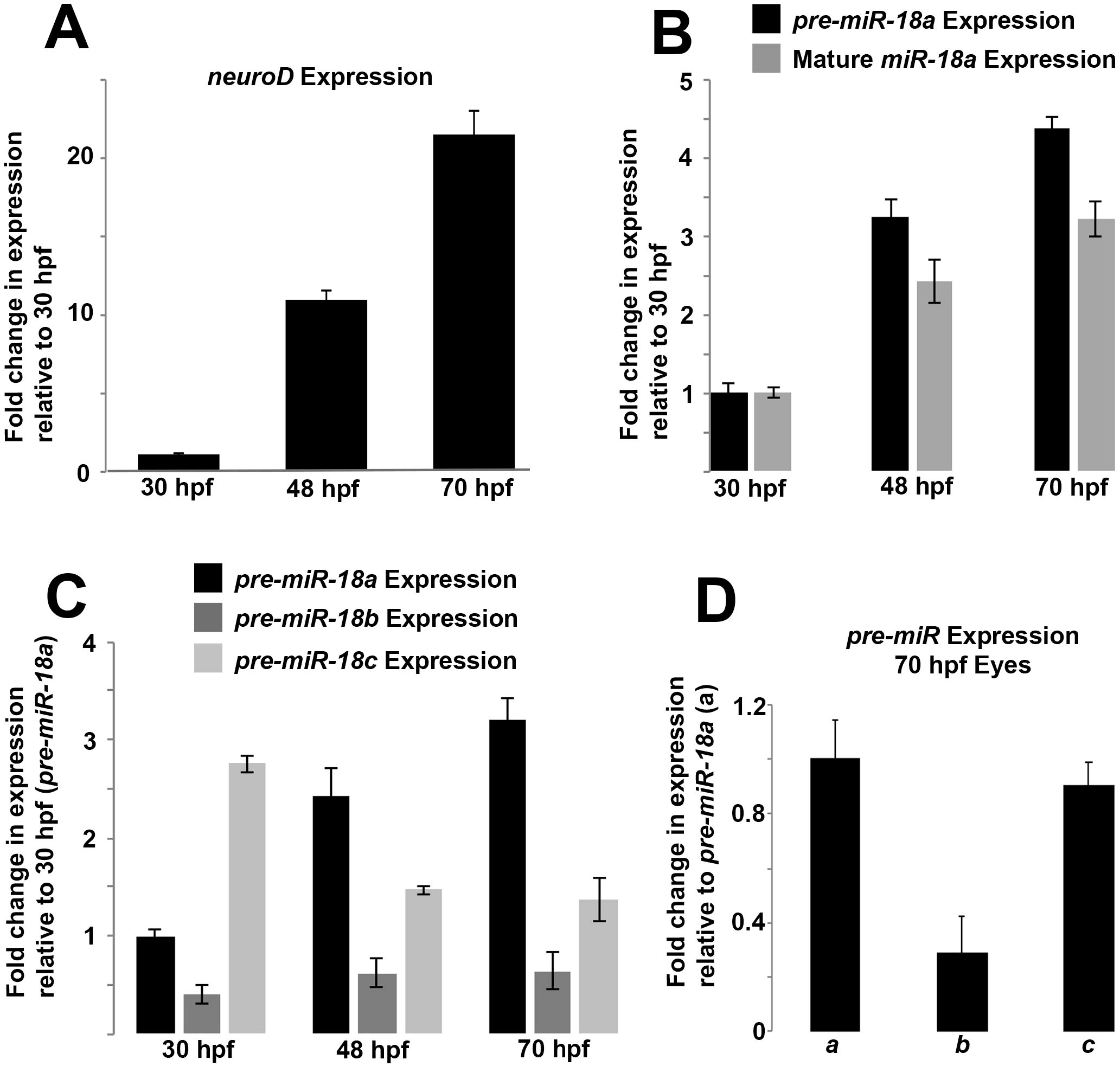
During development, *pre-miR-18a* expression increases proportionally along with *neuroD* mRNA and mature *miR-18a*. (A) *neuroD* mRNA expression in the developing brain and retina between 30 and 70 hpf. (B) Comparison between *pre-miR-18a* and mature *miR-18a* expression changes in the developing brain and retina between 30 and 70 hpf. (C) Comparison between *pre-miR-18a, pre-miR-18b* and *pre-miR-18c* expression changes in the developing brain and retina between 30 and 70 hpf. (D) Comparison between expression of *pre-miR-18a* (a), *pre-miR-18b* (b) and *pre-miR-18* (c) in the eyes only at 70 hpf. Error bars represent standard deviation; n=40 whole heads (A-C) or 40 whole eyes (D) from AB WT embryos per sample.

### Morpholino-induced knockdown of *miR-18a, miR-18b* or *miR-18c* increases the number of mature photoreceptors at 70 hpf

To determine if *miR-18* miRNAs regulate photoreceptor development, morpholino oligonucleotides targeted to the mature sequences of *miR-18a*, *mir-18b* or *miR-18c* were injected into embryos at the single cell stage. Compared with embryos injected with standard control morpholinos, morpholinos targeting each of the three *miR-18* types resulted in a greater number of cone photoreceptors at 70 hpf (Figure 2A,C); *miR-18a* knockdown was verified by in-situ hybridization (Figure 2B). None of the three morpholinos altered the number of BrdU+ cells (Figure 2D), indicating that the *miR-18* miRNAs regulate photoreceptor differentiation, but do not regulate the cell cycle. The high degree of sequence similarity between the three *miR-18* types and the identical effect of morpholinos targeted to each suggest that each morpholino might comprehensively knock down *miR-18a*, *b* and *c.* This potential cross-reactivity makes it difficult to determine which *miR-18* is most important regulator of photoreceptor development, but the results indicate that among post-mitotic cells of the photoreceptor lineage, *miR-18* miRNAs regulate differentiation.

**Figure 2.**
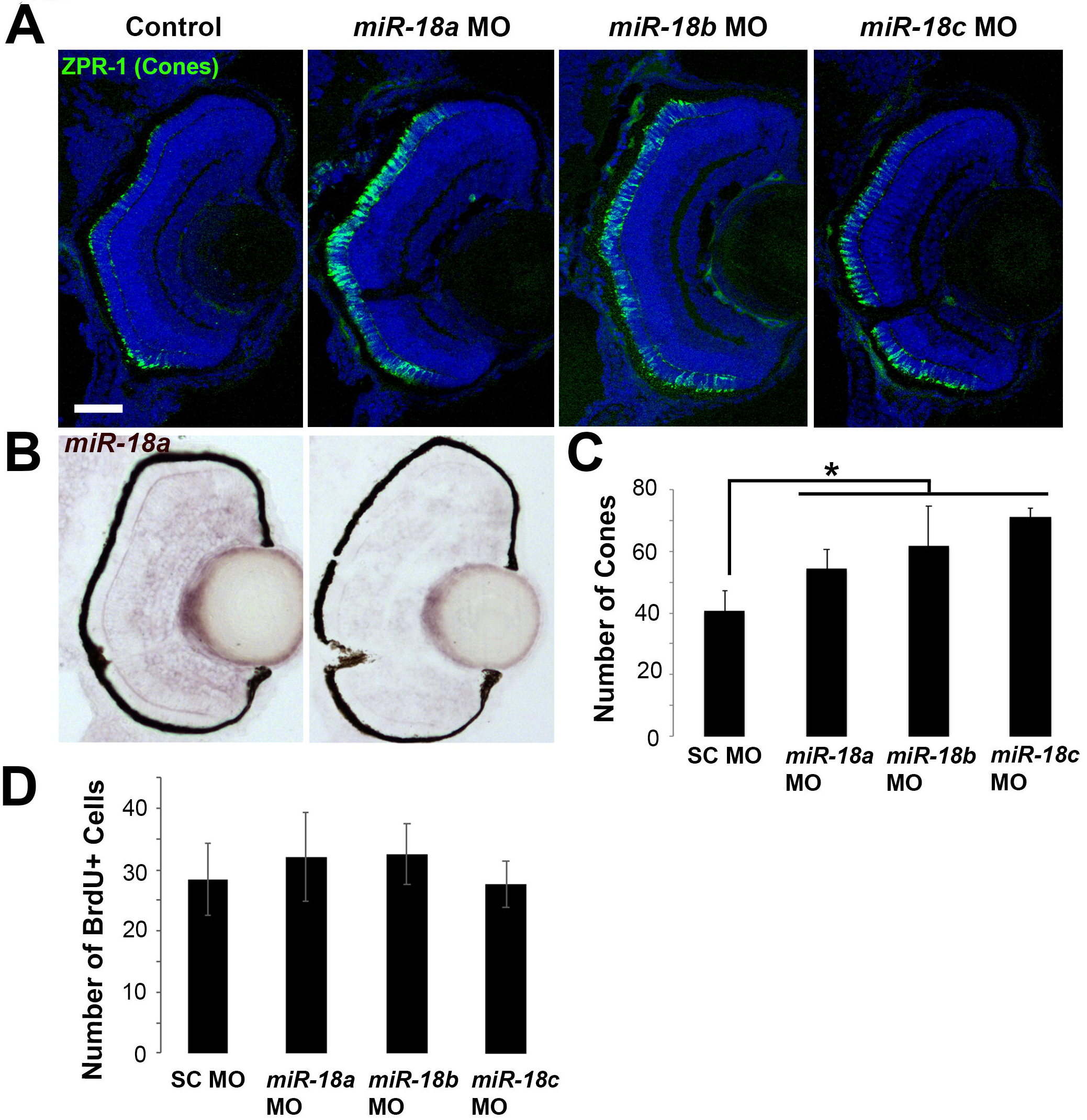
Knockdown with morpholinos targeted to *miR-18a*, *miR-18b* or *miR-18c* results in more differentiated cone photoreceptors. (A) ZPR-1 (cone) immunolabeling in 70 hpf larvae that were injected at the single-cell stage with standard control, *miR-18a, miR-18b* or *miR-18c* morpholinos. (B) Cone photoreceptor counts presented as the mean of one eye per fish (n=3) counted in the centermost cross-section in the vicinity of the optic nerve. (C) In situ hybridization for *miR-18a*, comparing expression in larvae injected with standard control morpholino (left) with *miR-18a* knockdown (right). (D) Total number of BrdU-labeled cells presented as the mean of one eye per fish (n=3) counted in the centermost cross-section in the vicinity of the optic nerve. Error bars show standard deviation and counts were statistically compared using a Student’s t-test.

### *miR-18a* directly interacts with *neuroD* mRNA

The similarity in the timing of expression between *pre-miR-18a*, mature *miR-18a* and *neuroD* suggests that, in the developing retina, *miR-18a* might regulate NeuroD. To determine if *miR-18a* directly interacts with *neuroD* mRNA, a dual luciferase assay was performed on HEK 293 cells transfected with pGL3-control firefly luciferase vector into which a 66 nt portion of the *neuroD* 3’ UTR was cloned immediately 3’ of the luciferase gene. The wild-type vector had the normal predicted target site for *miR-18a* on the *neuroD* 3’ UTR (CACCTTA) and a negative control vector was created with this predicted target site mutated to TTTTTTT (Figure 3A). As an internal transfection control, cells were cotransfected with pRL-TK vector with constitutive expression of *Renilla* luciferase. Following cell culture and transfection, cell lysates were treated with either *miR-18a* mimic or a negative control miRNA mimic (*let-7a*) not predicted to interact with the *neuroD* 3’ UTR (Figure 3B). Binding of the miRNA to the cloned *neuroD* sequence was predicted to suppress the level of firefly luciferase expression relative to *Renilla* luciferase. In the wild-type *neuroD* vector relative to negative controls, *miR-18a* resulted in a significant decrease in firefly luciferase expression (Figure 3C). In the mutated *neuroD* vector, relative to negative controls, *miR-18a* did not significantly affect firefly luciferase expression. These results indicate that *miR-18a* binds to the 3’ UTR of *neuroD* at the predicted target site and suggests that *miR-18a* functions to negatively regulate NeuroD translation.

**Figure 3.**
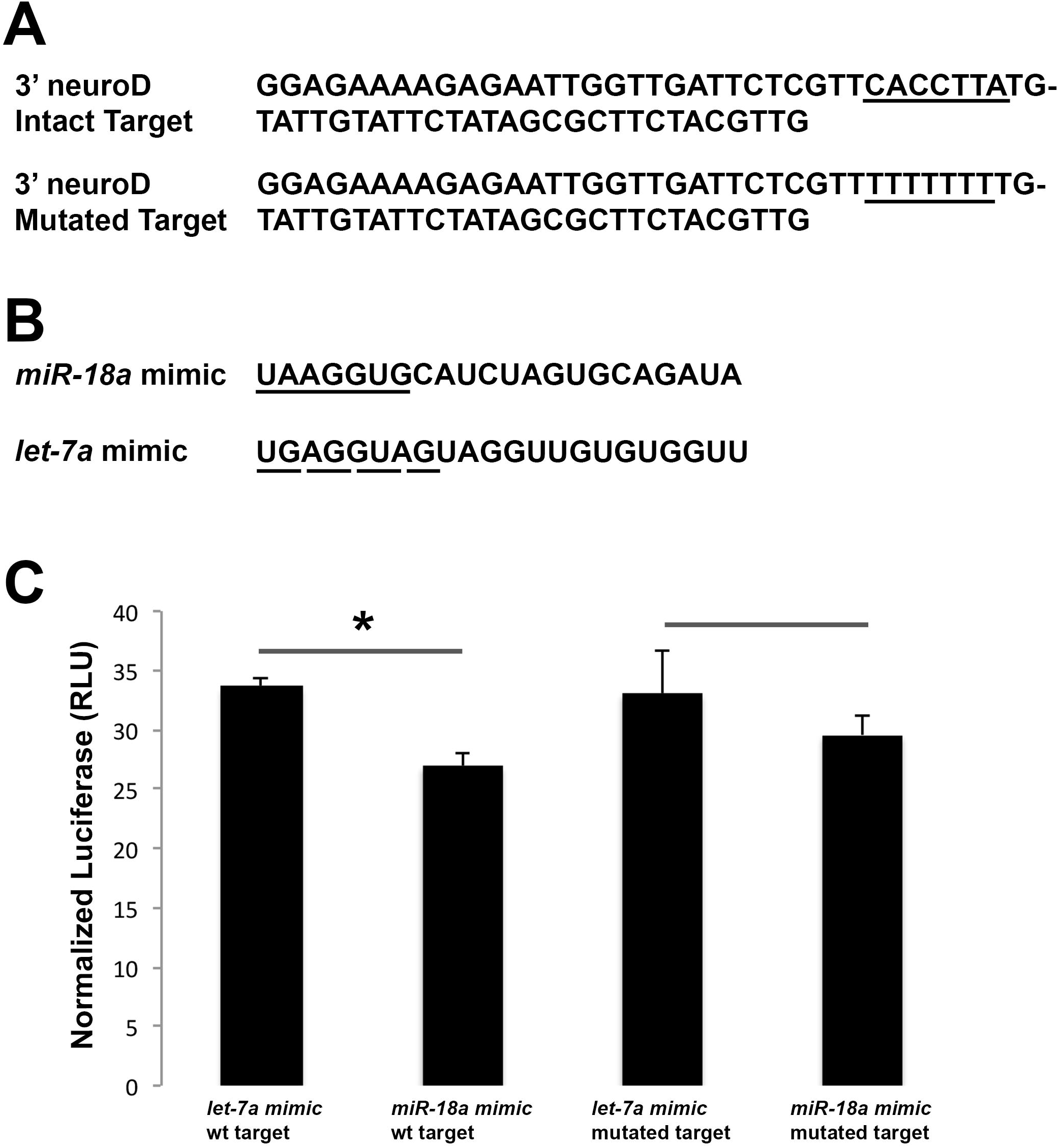
Dual luciferase assay showing direct interaction between miR-18a and its predicted target site in the 3’ UTR of neuroD mRNA. (A) The 66 bp portion of the *neuroD* 3’ UTR sequence inserted into the pGL3 vector with the predicted intact (top) or mutated (bottom) *miR-18a* target site underlined. (B) Sequences for the *miR-18a* and *let-7a* (negative control) mimics that were co-transfected into cells. The seed sequence of the *miR-18a* mimic is underlined with a solid line, which is complementary to the underlined target sequence in (A); the seed sequence of the *let-7a* mimic, used as a negative control, is underlined with a dashed line and is not complementary to any portion of the *neuroD* 3’ UTR sequence in (A). (C) Firefly luciferase from the pGL3 vector shown relative to constitutive firefly luciferase from the pRL-TK vector, compared between treatments with the *let-7a* mimic (negative control) and *miR-18a* mimic and shown for both the intact predicted *miR-18a* target site and the mutated target site. Error bars show standard deviation; values for *let-7a* and *miR-18a* mimic were compared with a Student’s t-test and asterisks indicate p<0.05.

### *miR-18a* mutants generated by CRISPR/CAS9 gene editing

To determine the role of *miR-18a*, independent of *miR-18b* or *c*, in regulating NeuroD and photoreceptor differentiation, CRISPR/Cas9 gene editing was used to generate a *miR-18a*^-^/^-^ mutant line. An appropriate CRISPR target site was not available within the 22bp sequence coding for the mature miRNA molecule, so a target site was chosen within the sequence for the larger precursor molecule (*pre-miR-18a*), and mutations here were predicted to interfere with processing by Dicer into the mature miRNA. This method produced animals with a 25 nt insertion within the sequence (Figure 4A) that normally produces the stem-loop precursor molecule (Figure 4B). This insertion introduced a restriction site for AleI that is not present in WT DNA (Figure 4A), and restriction analysis was subsequently used for genotyping (Figure 4C). To produce a stable mutant line, F2 generation heterozogous fish were in-crossed to produce homozygous mutants, and F3 generation homozygous mutants were in-crossed to produce homozygous mutant embryos. TaqMan qPCR, specific for the mature 22 nt *miR-18a*, was then used to compare expression of mature *miR-18a* in mutant and wild type fish. In these mutants compared with wild-type fish at 70 hpf, mature *miR-18a* expression was reduced by more than 14-fold (Figure 4D), demonstrating that *miR-18a* mutants lack mature *miR-18a*.

**Figure 4.**
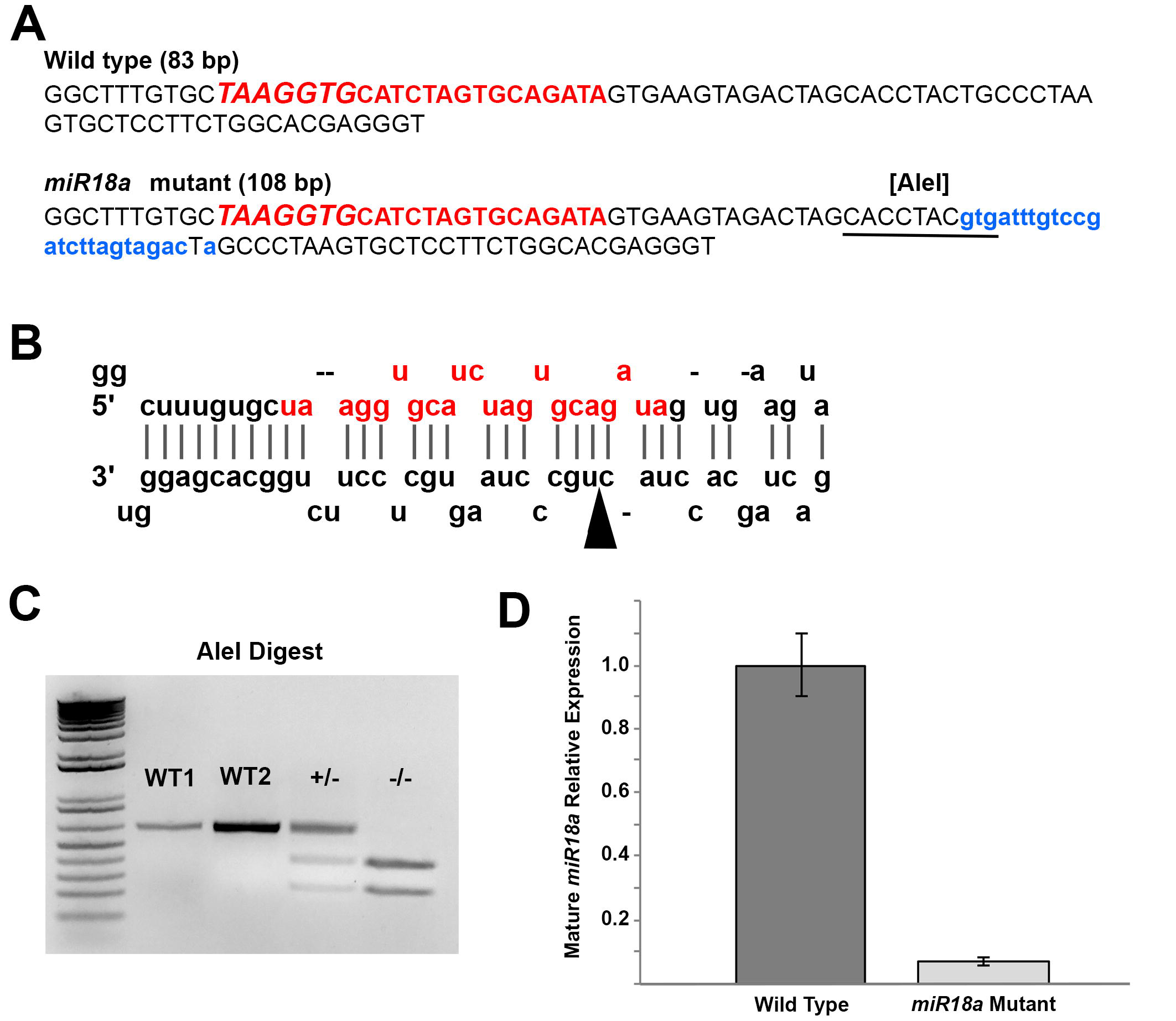
Characterization of the *miR-18a*^-/-^ mutant line. (A) Comparison between WT and mutant genomic sequences corresponding to *pre-miR-18a* and in the mutant, the 25 bp insertion is shown in lowercase, blue lettering and the introduced Alel restriction cut site is underlined. The uppercase red lettering shows the genomic sequence corresponding to mature *miR-18a*, with the seed sequence in larger italics. (B) The predicted stem loop arrangement for the WT *pre-miR-18a* RNA molecule with the mature *miR-18a* sequence shown in red (adapted from www.mirbase.org); in the mutant sequence, the 2 5-base insertion location is indicated by the arrowhead. (C) Genotyping of *miR-18a* mutants using Alel restriction digest. In WT fish, the 626 bp PCR product remains uncut, with clear intensity distinctions between 200 ng (Wl) and 400 ng (WT2) PCR product. In heterozygous mutants (+/−) 50% (~200 ng) of the PCR product is cut into smaller fragments and in homozygous mutants (-/-) 100% (400 ng) of the PCR product is cut into smaller fragments. (D) TaqMan qPCR showing, compared with WT at 70 hpf, the relative absence of mature *miR-18a* in mutant fish. Error bars represent standard deviation, n=40 embryo heads.

### *miR-18a* regulates NeuroD and the timing of photoreceptor differentiation

To determine if *miR-18a* regulates NeuroD, Western blot was used to compare NeuroD protein levels in 48 hpf embryos (heads) between fish injected with standard control or *miR-18a* morpholinos, and between WT and *miR-18a* mutant fish. Knockdown of *miR-18a* resulted in a 32% increase in NeuroD protein and *miR-18a* mutation resulted in a 20% increase in NeuroD protein indicating that in the developing brain and retina, *miR-18a* suppresses the level of NeuroD (Figure 5A,B). These data also indicate that broader knockdown of *miR-18*(a, b and c) with morpholinos may have a greater effect on NeuroD protein levels than *miR-18a* mutation, where *miR-18b* and *c* are still functional. Then, to determine if *miR-18a*, independent of *miR-18b* or *c*, regulates photoreceptor differentiation, the numbers of mature photoreceptors were compared between WT and *miR-18a* mutant fish. Immunohistochemistry for the red/green cone marker Arrestin-3a was used to label a subset of cone photoreceptors, while *in-situ* hybridization for the mature rod marker *rhodopsin* was used to label rods. Larvae were placed in lOmM BrdU solution for 20 minutes prior to sacrifice at 70 hpf. The *miR-18a* mutation resulted in a significantly greater number of both mature red/green cones and rods, whereas the numbers of BrdU-labeled cells remained invariant (Figure 5C-F). This indicates that within the photoreceptor lineage, *miR-18a* regulates photoreceptor differentiation but does not regulate the cell cycle. By 6 dpf, the numbers of mature photoreceptors do not differ between mutant and wild-type fish (not shown), indicating that *miR-18a* does not regulate cell fate or the total numbers of photoreceptors generated. Taken together, these results indicate that, among post-mitotic cells already determined to become photoreceptors, *miR-18a* functions to regulate the timing of photoreceptor differentiation.

**Figure 5.**
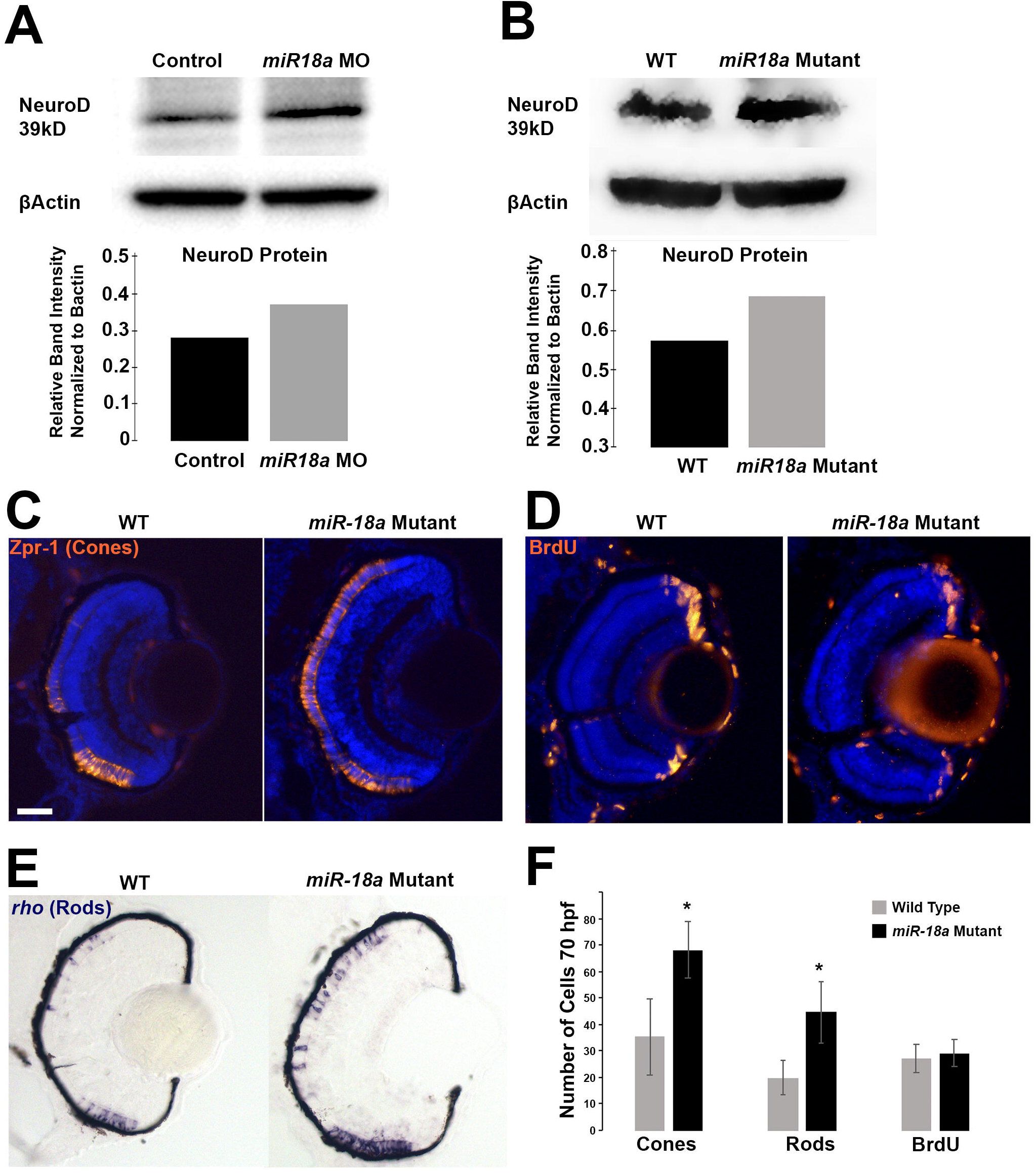
Loss of *miR-18a* increases NeuroD protein levels and the number of differentiated photoreceptors. Western blot on 48 hpf embryo heads (n=40) comparing NeuroD protein levels between standard control MO injected and *miR-18a* MO injected embryos (A) and between WT and *miR-18a^-/-^* mutant embryos (B) with corresponding quantification graphs. In WT compared with *miR-18a* mutant larvae at 70 hpf, immunolabeling for mature cone photoreceptors (C: Zpr-1) and cells in S-phase of the cell cycle (D: BrdU); and in-situ hybridization for rod photoreceptors (E: *rhodopsin*); scale bar =0.50 μm. (F) Quantification of cones (n≥14 larvae), rods (n≥8 larvae) and BrdU+ cells (n≥7 larvae). Error bars represent standard deviation; cell counts compared with a student’s t-test and asterisks indicate p<0.05.

### Increased *miR-18a* expression suppresses NeuroD protein levels and photoreceptor differentiation

TGIF1 is a transcriptional repressor in the TGFβ signaling pathway (Lenkowski et al., 2013) and, compared with WT larvae at 70 hpl, *tgif1* mutant larvae were observed to have fewer differentiated cone photoreceptors (Figure 6A,B). *miR-18a* was investigated as the possible mediator of this phenotype and, in 70 hpf *tgif1* mutants compared with WT, in-situ hybridization indicated increased retinal expression of *miR-18a* (Figure 6A,B) while qPCR showed higher levels of *pre-miR-18a* expression (Figure 6C). To determine if increased *miR-18a* expression mediates the loss-of-cone phenotype in *tgif1* mutants, morpholinos were used to knock down *miR-18a* in *tgif1* mutant larvae. Compared with standard control morpholino-injected larvae (SC MO) at 70 hpf, *mir-18a* knockdown in *tgif1* mutants fully rescued the deficiency in cone differentiation (Figure 6A,B), indicating that the lack of cone phenotype is mediated through *miR-18a.* To determine if, in the *tgif1* mutant retina, *miR-18a* post-transcriptionally regulates NeuroD, qPCR and Western blot were used to compare mRNA expression and NeuroD protein, respectively, between WT and *tgif1* mutant larvae. Compared with WT at 70 hpf, *tgif1* mutants have identical levels of *neuroD* expression (Figure 6D) but reduced NeuroD protein (Figure 6E,F). These results show that, in *tgif1* mutants, NeuroD protein levels are suppressed at the post-transcriptional level and suggest that this regulation is mediated through increased levels of *miR-18a*.

**Figure 6.**
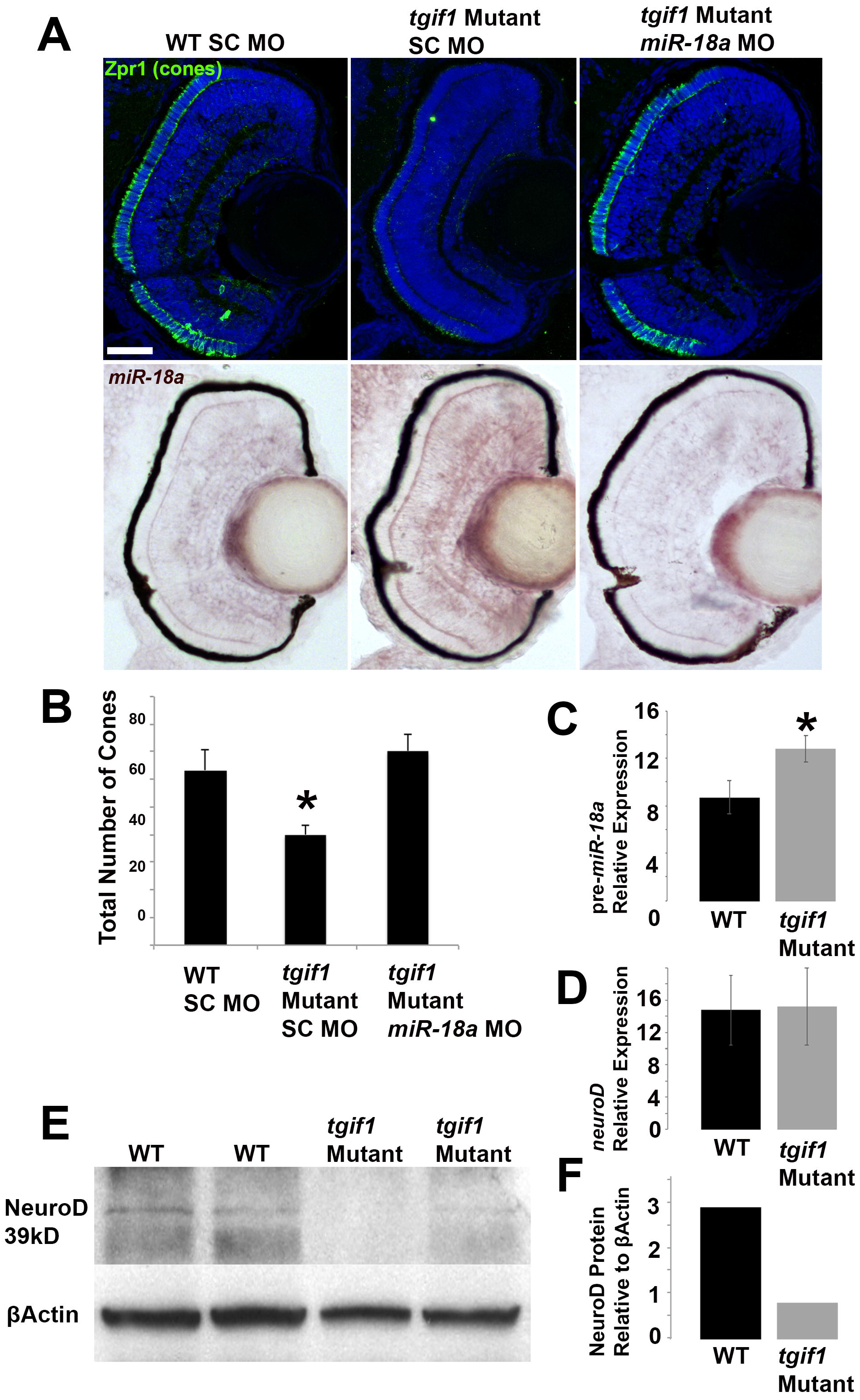
*tgif1* mutant larvae fewer photoreceptors, increased *miR-18a* expression and reduced NeuroD protein levels. (A) Immunolabeling for cone photoreceptors (Zpr-1) at 70 hpf in WT larvae injected with standard control morpholinos (SC MO), *tgif1* mutant larvae injected with SC MO, *tgif1* mutant larvae injected with *miR-18a* morpholinos, with corresponding images showing in-situ hybridization for *miR-18a* in the same retinas; scale bar = 50 μm. (B) Cone photoreceptor counts in retinal cross-sections (n=3 larvae each) in fish corresponding to the images in (A). (C) Standard qPCR showing *pre-miR-18a* expression in 70 hpf WT larvae compared with *tgif1* mutants (n=40 heads); normalized to *pctin* and shown relative to *let-7b* expression. (D) Standard qPCR comparing *neuroD* mRNA expression in 70 hpf larvae between WT and *tgif1* mutants (n=40 heads); normalized to *βactin* and shown relative to *ccnb1* expression. (E) Western blot showing NeuroD protein levels in 70 hpf WT compared with *tgif1* mutant fish (n=40 heads). (F) Quantification of the average band intensities in E. All error bars represent standard deviation and comparisons were made with Student’s t-tests (asterisks indicate p<0.05).

## DISCUSSION

Generating the correct type and number of neurons in the developing brain requires precise temporal and spatial regulation of mechanisms that specify progenitor cell fate, determine the timing of cell cycle exit and regulate differentiation (recent reviews: Cepko, 2014; Mattar & Cayouette, 2015; Stenkamp, 2015; Wang & Cepko, 2016). This control is partially accomplished through transcription factors that regulate mRNA expression levels (reviewed in J. A. Brzezinski & Reh, 2015; Gregory-Evans, Wallace, & Gregory-Evans, 2013). NeuroD is a bHLH transcription factor that, within the photoreceptor lineage, governs the cell cycle through Delta-Notch signaling and, in newly generated photoreceptors, governs differentiation through a separate mechanism (Taylor et al., 2015). Little is currently known about how NeuroD is regulated, but there is evidence that in the retina of Medaka fish the transcription factor Six6 influences *neuroD* expression (Conte et al., 2010), and Insm1a functions upstream of NeuroD in zebrafish (Forbes-Osborne, Wilson, & Morris, 2013). Here we identify a mechanism through which *miR-18a* post-transcriptionally regulates NeuroD protein levels and the timing of photoreceptor differentiation. We propose that the fine-tuning of NeuroD protein levels by *miR-18a* provides a mechanism through which NeuroD differentially governs both cell cycle exit in photoreceptor progenitors and the timing photoreceptor differentiation.

Within the photoreceptor lineage, NeuroD governs two sequential events—cell cycle exit and differentiation—through distinct mechanisms (Taylor et al., 2015). Retinal expression of *neuroD* mRNA increases steadily between 30 and 70 hpf, throughout the retinal neuroepithelium by 38 hpf and in all developing photoreceptors in the ONL by 48 hpf (Malgorzata J. Ochocinska & Hitchcock, 2007). Despite this relative uniform expression of *neuroD* among photoreceptor progenitors, the spatiotemporal pattern of events governed by NeuroD are complex. Among photoreceptor progenitors, NeuroD governs the cell cycle through intercellular Delta-Notch signaling, with the first photoreceptor progenitors beginning to exit the cell cycle around 48 hpf (Stenkamp, 2007). At this time, *neuroD* mRNA is expressed in all ONL cells (Malgorzata J. Ochocinska & Hitchcock, 2007), but photoreceptor genesis begins in only a small ventronasal patch around this time (Schmitt & Dowling, 1999). Photoreceptor genesis then spreads peripherally until most ONL cells have exited the cell cycle by about 60 hpf (Stenkamp, 2007). Photoreceptor differentiation, coincident with expression of the mature photoreceptor markers Rhodopsin (rods) and Arrestin3a (red/green cones), is also governed by NeuroD but lags slightly behind cell cycle exit and is completed by 72 hpf (Stenkamp, 2007). This clear and tightly controlled spatiotemporal pattern of photoreceptor genesis and differentiation, despite the uniform expression of *neuroD* among photoreceptor progenitors, suggests that post-transcriptional mechanisms could regulate NeuroD function.

MicroRNAs post-transcriptionally regulate protein levels by binding to the 3’UTR of target mRNA and blocking translation (Djuranovic, Nahvi, & Green, 2012; Iwakawa & Tomari, 2015; Valencia-Sanchez, Liu, Hannon, & Parker, 2006; Zeng, Yi, & Cullen, 2003) and in some cases causing mRNA degradation (Huntzinger & Izaurralde, 2011). In the developing CNS, miRNAs regulate stem and progenitor cell proliferation, cell fate specification and neural differentiation (Pham & Gallicano, 2012; Shi et al., 2010), as well as the timing of retinal neurogenesis (La Torre et al., 2013). The microRNAs *miR-18a*, *miR-18b* and *miR-18c* share a 7-base seed region that is predicted to interact with the 3’ UTR of *neuroD* and, if expressed in the retina, could post-transcriptionally regulate NeuroD and photoreceptor genesis. The genes for these three *miR-18* molecules are on different chromosomes, and their expression is, therefore, presumably controlled by different regulatory mechanisms.

MicroRNAs are initially expressed as long primary transcripts and then cleaved into smaller pre-miRNAs in the nucleus by the Drosha enzyme complex (Zeng et al., 2005). Then, in the cytoplasm, pre-miRNAs are cleaved again and processed into single-stranded mature miRNAs by the RNAse DICER and the RISC loading complex (Winter et al., 2009). MicroRNA biogenesis is a complex process but, for many miRNAs, precursor and mature miRNA expression levels are closely correlated (Nepal et al., 2016; Powrózek, Mlak, Dziedzic, Małecka-Massalska, & Sagan, 2018). Accordingly, we show that in the developing brain and retina, *pre-miR-18a* expression increases proportionally with, and can serve as an accurate proxy for, mature *miR-18a* expression. This is an advantage because, compared with mature miRNAs that must be amplified with special qPCR kits (e.g. TaqMan) that might not fully discriminate between nearly identical miRNAs (e.g. *miR-18a*, *b*, *c*), the longer pre-miRNAs have unique sequences and can be easily discriminated using standard qPCR. Taking advantage of this, we show that each of the precursor molecules for *miR-18a*, *miR-18b* and *miR-18c* are all expressed in the developing brain and retina, but the timing of their expression differs, which might be key to their functions. Most photoreceptor genesis occurs between 48 and 72 hpf and, like *neuroD, pre-miR-18a* and mature *miR-18a* expression increase steadily between 30 and 70 hpf. In comparison, *pre-miR-18c* expression peaks at 30 hpf and then is rapidly downregulated, while *pre-miR-18b* expression remains substantially lower (than *pre-miR-18a* or c) throughout embryonic development. These data indicate that, in the brain and retina, *miR-18a, b* and *c* function during distinct developmental time frames, but *miR-18a* expression most closely correlates with the timing of *neuroD* expression and photoreceptor genesis.

Even though expression data suggest that *miR-18a*, *b* and *c* function during different developmental events, morpholinos targeted to each of these miRNAs result in an identical phenotype in which 70 hpf larvae have significantly more mature photoreceptors. This suggests that, due to their nearly identical sequences, each morpholino knocks down all three miRNAs and these miRNAs may have overlapping functions. Redundancy among miRNAs occurs commonly and may be important for cooperative translational repression (Fischer, Handrick, Aschrafi, & Otte, 2015). Knockdown of multiple, redundant miRNAs by a single morpholino has been documented for other miRNA groups (Alex S. Flynt et al., 2009), indicating that morpholino oligonucleotides can be an effective tool for understanding cooperative function of multiple miRNAs (Alex Sutton Flynt et al., 2017). In contrast, removing individual miRNAs using gene editing (e.g. CRISPR/Cas9) or knockout techniques sometimes does not produce a phenotype (Olive, Minella, & He, 2015), because redundant miRNAs might partially or fully compensate for functional loss of a single miRNA (Bao et al., 2012; Gurtan & Sharp, 2013; Ventura et al., 2008). This redundancy, however, does not indicate that familial miRNAs are merely functional replicates of one another. Expression regulation and feedback mechanisms under different circumstances can confer distinct roles for what are considered redundant miRNAs (Olive et al., 2015). Accordingly, differential expression of *pre-miR-18a*, *b* and *c* in the brain and retina suggest that these miRNAs may have functional specializations during distinct developmental events. Based on the overlap in expression between *miR-18a* and *neuroD, miR-18a* was investigated, independent of *miR-18b* or *c*, as a potential regulator of NeuroD and photoreceptor genesis. First, an *in vitro* double luciferase assay showed that *miR-18a* suppresses translation through direct interaction with the 3’UTR of *neuroD*, indicating that *miR-18a* post-transcriptionally regulates NeuroD. This is consistent with the widely demonstrated roles of miRNAs to suppress translation through direct interaction with the 3’UTR region of target mRNAs (Humphreys, Westman, Martin, & Preiss, 2005; Valencia-Sanchez et al., 2006; van den Berg, Mols, & Han, 2008; Zeng et al., 2003). Then, to determine if *miR-18a* regulates NeuroD and photoreceptor differentiation, CRISPR-Cas9 gene editing was used to generate a stable mutant line that lacks mature *miR-18a.* In *miR-18a* morphants and mutants, at 48 hpf when photoreceptor differentiation begins, Western blot showed that NeuroD protein levels are increased by 32% and 20%, respectively. This indicates that, in the wild-type retina during the time of photoreceptor differentiation, *miR-18a* suppresses NeuroD protein levels. This also suggests that concurrent knockdown of *miR-18a, b* and *c* by the *miR-18a* morpholino has a greater effect on NeuroD protein than mutation of *miR-18a* alone. Finally, identical to *miR-18*(*a, b* and c) morphants, 70 hpf *miR-18a* mutant larvae have significantly more mature photoreceptors, whereas the numbers of cells in the cell cycle are equivalent. By 6 days post-fertilization, after embryonic retinal development is complete, the total number of mature rods and cones does not differ between *miR-18a* mutant and WT fish. Taken together, these data indicate that, among post-mitotic cells within the photoreceptor lineage, *miR-18a* regulates the timing of photoreceptor differentiation. This is consistent with *miR-18a* functioning through NeuroD, which also does not regulate photoreceptor fate but, within the photoreceptor lineage, governs differentiation (M. J. Ochocinska & Hitchcock, 2009; Taylor et al., 2015). These results also demonstrate that while, based on their sequences, *miR-18a*, *b* and *c* may be considered functionally redundant, *miR-18b* and *c* do not fully compensate for loss of *miR-18a*.

*Tgif1* mutant larvae were observed to have increased expression of *miR-18a* throughout the retina, providing an opportunity to determine the effects of *miR-18a* gain-of-function. In *tgif1* mutants compared with WT, despite equivalent *neuroD* mRNA expression, NeuroD protein levels are lower and there are fewer differentiated photoreceptors. Knockdown of *miR-18a* in *tgif1* mutants fully rescues the photoreceptor deficiency, indicating that the elevated *miR-18a* expression in these mutants can account for the absence of differentiated photoreceptors. The TGIF1 protein is a transcriptional corepressor in the TGFβ pathway that, in the adult zebrafish retina, is critical for injury-induced photoreceptor regeneration (Lenkowski et al., 2013). The role of the TGFβ pathway has not been investigated during embryonic retinal development, but our results suggest a role in regulating miRNAs and photoreceptor differentiation and that *miR-18a* functions downstream of Tgif1. These data show that *miR-18a* gain-of-function in the *tgif1* mutants produces a phenotype opposite that of the *miR-18a* morphants or mutants and, taken together, show that *miR-18a* post-transcriptionally regulates NeuroD and, thereby, governs photoreceptor differentiation.

In mutant or morphant fish lacking *miR-18a*, photoreceptors differentiate at a faster rate without any obvious defects in retinal morphology, suggesting that *miR-18a* inhibition could have therapeutic potential. Recent studies demonstrate regenerative potential in the mouse retina, in which EGF stimulates Müller glia to proliferate (Ueki & Reh, 2013) and Ascl1a confers permissive reprogramming in Müller glia that generate neuronal progenitor cells (Pollak et al., 2013). Few of these Müller glia-derived progenitors differentiate into photoreceptors, but this problem might be overcome through creating a therapeutic environment that favors photoreceptor differentiation (e.g. through *miR-18a* inhibition). This could be accomplished using RNA silencing or CRISPR/Cas9 gene editing, both of which have been successfully employed *in vivo* to knock down miRNAs (Chang et al., 2016; Shah, Ferrajoli, Sood, Lopez-Berestein, & Calin, 2016). Therapeutic approaches using these methods are becoming more feasible as improvements are made in molecule delivery to target tissues using viral vectors (Zhu et al., 2017) and in creating CRISPR tools that can be activated *in vivo* (Dow et al., 2015; Hirosawa et al., 2017). Using viral vectors, CRISPR-based cellular reprogramming was recently shown to prevent photoreceptor degeneration in a mouse model for the retinal disease retinitis pigmentosa (Zhu et al., 2017).

In conclusion, the data presented here demonstrate that during normal retinal development, *miR-18a* regulates the timing of photoreceptor differentiation, and indicate that *miR-18a* functions through post-transcriptional regulation of NeuroD protein levels.

This is consistent with the known role of NeuroD in governing differentiation in postmitotic photoreceptors (Taylor et al., 2015) and the functions of some miRNAs to effectively uncouple transcription and translation in order to ensure the correct spatiotemporal expression of proteins (Bao et al., 2016; McLaughlin, Smith, Catrina, & Bratu, 2018; Parchem et al., 2015). We propose that within the photoreceptor lineage, following cell cycle exit, fine-tuning of NeuroD protein levels by *miR-18a* regulates the spatiotemporal pattern of photoreceptor differentiation. The importance of the spatiotemporal pattern of photoreceptor genesis is not yet understood, but could affect development of the correct distribution, positioning and, in zebrafish, the rigid spatial mosaic of cone photoreceptors (Allison et al., 2010; Raymond & Barthel, 2004; Raymond et al., 2014) that are essential for normal visual function.

## AUTHOR CONTRIBUTIONS

SMT performed this research at the University of Michigan and the University of West Florida. EG and PM performed this research at the University of West Florida. Writing and editing were performed by SMT, EG and PFH.

## ACKNOWLEDGEMENTS

This work was supported by grants from the National Institutes of Health (NEI)-R01EY07060 (PFH), T32EY013934 (SMT), P30EY07003 (PFH) and an unrestricted grant from the Research to Prevent Blindness, New York. The authors declare no competing financial interests. The authors thank Laura Kakuk-Atkins and Dilip Pawar for technical assistance. Fish lines and reagents provided by ZIRC were supported by NIH-NCRR Grant P40 RR01.

